# Methods for the generation of heritable germline mutations in the disease vector *Culex quinquefasciatus* using CRISPR/Cas9

**DOI:** 10.1101/609206

**Authors:** Ming Li, Ting Li, Nannan Liu, Robyn Raban, Xuegui Wang, Omar S. Akbari

## Abstract

*Culex quinquefasciatus* is vector of many diseases that adversely impact human and animal health; however, compared to other mosquito vectors limited genome engineering technologies have been characterized for this vector. CRISPR-Cas9 based technologies are a powerful tool for genome engineering and functional genomics and consequently have transformed genomics studies in many organisms. Our objective was to improve upon the limited technologies available for genome editing in *Cx. quinquefasciatus* to create a reproducible and straightforward method for CRISPR-Cas9-targeted mutagenesis in this vector. Here we describe methods to both improve embryo survival rates as well as mutagenesis rates by optimizing injection supplies and equipment, embryo injection procedures, embryo handling and gRNA target design. Through these efforts, we achieved embryo survival rates and germline mutagenesis rates that greatly exceed any previously reported rates in this vector. This work was also the first characterize the *white* gene marker, which is a valuable phenotypic marker for future transgenesis or mutagenesis of this vector. In the end, these tools provide the framework for future functional genomic studies in this important disease vector and may support the development of future gene drive and genetic technologies that can be used to control this vector.

## Introduction

*Culex quinquefasciatus* is a primary vector of West Nile virus (WNV), eastern equine encephalitis virus, Saint Louis encephalitis virus, lymphatic filariasis and avian malaria (Reisen et al. 2005; Jones et al. 2002; Petersen & Roehrig 2001; LaPointe et al. 2005). Since the introduction of WNV to the United States in 1999, there have been over 48,000 confirmed cases of WNV in the United States (CDC, ArboNET). While this is certainly and underestimate of the overall impact of this disease in the US, the global impact of lymphatic filariasis is far greater. Current estimates indicate that despite remarkable mass drug administration (MDA) programs, millions of people are currently infected with lymphatic filariasis, which is considered to be one of the leading global causes of disability (Gyapong et al. 2018). Furthermore, there are many areas of the world where MDA is not expected to eradicate this disease, but mosquito control could be the key to eliminating this disease (Koudou et al. 2018). Moreover, *Cx. quinquefasciatus* has established itself as the dominant vector of avian malaria in many island habitats, and in some instances caused the extinction of many rare bird species (LaPointe et al. 2012). Thus, technologies to improve *Cx. quinquefasciatus* control are essential to reduce and potentially eliminate these diseases. However, the widespread development of insecticide resistance in *Cx. quinquefasciatus*, the primary method for control of this vector, has made it imperative that alternative control methods are developed for this vector.

Genome engineering technologies facilitate important functional genomic research as well as the development of tools required to create genetic control strategies for important disease vectors. Recently, the clustered regularly interspaced short palindrome repeats (CRISPR)-associated protein 9 (CRISPR-Cas9) system has been used to generate somatic and heritable germline mutations and genetic drive technologies in *Aedes* (Li, Bui, et al. 2017; Kistler et al. 2015) and *Anopheles* (Li et al. 2018; Dong et al. 2018; Gantz et al. 2015; Hammond et al. 2016; Kyrou et al. 2018) disease vectors. In this system, a Cas9 endonuclease and a small guide RNA (sgRNA) complementary to the target site facilitate site directed double stranded genome breaks, which are repaired by homology directed repair (HDR) or nonhomologous end joining (NHEJ). Despite this being a useful mutagenesis tool in other vector species, there has been limited emphasis on improving CRISPR genome engineering technologies to study and develop new control tools for *Culex* disease vectors. In 2016, transcription activator-like effector nucleases (TALEN) and CRISPR-Cas9 genome engineering technologies were used to generate frameshift mutations to disrupt the function of an insecticide resistance gene in *Cx. quinquefasciatus* (Itokawa et al. 2016), but no additional CRISPR/Cas9 mutagenesis work has been published for this vector. Furthermore, with the publication of the *Cx. quinquefasciatus* genome in 2010 (Arensburger et al. 2010), there have been few functional genomic studies of this species, which is unfortunate due to not only its importance a disease vector, but its unique biological characteristics (Severson & Behura 2012) including its susceptibility to diverse pathogens (i.e viral, nematode and protozoan) (Bartholomay et al. 2010), opportunistic blood feeding behavior (i.e. birds, humans and other mammals) as well as diverse geographic and habitat preferences. Therefore, in order to make CRISPR technologies more accessible to the research community, we aim to meticulously describe the development of a CRISPR-Cas9 mutagenesis system for *Cx. quinquefasciatus* targeting the *white* gene. This is the first characterization of this gene in *Cx. quinquefasciatus*, which has been an important phenotypic marker for genome engineering of other mosquito vectors (Coates et al. 1997). We hope that this tool will be useful for further functional genomics studies in this important disease vector and may lay the framework for the development of genetic control tools.

## Results

### Development of an CRISPR/Cas9 embryo microinjection protocol

We established efficient techniques for egg collection, pre-blastoderm stage embryo microinjection, and subsequent rearing and genetics. In brief, we first optimized the injection protocol by evaluating different types of capillary glass needles (quartz, aluminosilicate, borosilicate). The needle pulling settings were also optimized to minimize breakage and clogging during the injection procedures, while still maximizing embryo survival. Table S2 shows the optimal needle pulling parameters for all 3 needle types, but we found the aluminosilicate needles to produce the highest embryo survival at the most affordable costs.

We then conducted experiments to optimize the mosquito handling procedures. The mosquito mating, blood feeding and oviposition procedures were varied slightly with no large resulting effect, but egg raft separation and handling was key to obtaining high embryo survival rates. Methods that optimized egg separation and handling are outlined in the materials and methods. Furthermore, careful removal of the halocarbon injection oil from the eggs with a paintbrush, was also key to ensuring high embryo survival rates. Once the eggs were hatched, screening was performed my standard methods.

### Identification of CRISPR/Cas9 target sites

To test the efficiency of our CRISPR/Cas9 based genome editing platform in *Cx. quinquefasciatus*, we targeted the *white* (*w*) gene (CPIJ005542), which codes a protein critical for eye pigment transport. In other species, biallelic mutations in the *w* gene disrupts production of dark eye pigmentation and generates an easily screenable unpigmented eye color (Li, Bui, et al. 2017; Li et al. 2018; Ren et al. 2014; Bassett et al. 2014; Xue et al. 2018). Consequently, we designed three single-guide RNAs (sgRNAs) targeting three conserved regions of the third exon of the *w* gene (Fig. 2A). Target site conservation was confirmed in the CpipJ2 assembly of the Johannesburg strain of *Cx. quinquefasciatus* (www.vectorbase.org). Off-target effects were evaluated with CHOPCHOP v2 software (Labun et al. 2016), CRISPRdirect (https://crispr.dbcls.jp/) and a local sgRNA Cas9 package (Xie et al. 2014).

### Mutagenesis of the *white* gene locus in *Culex quinquefasciatus*

Our previous transgenesis work in *Aedes* (Li, Bui, et al. 2017) and *Anopheles* (Li et al. 2018) and our work the parasitoid wasp, *Nasonia vitripennis* (Li, Au, et al. 2017), demonstrated that sgRNA/Cas9 directed mutagenesis is dose dependent. Therefore, since *Cx. quinquefasciatus* eggs are larger than *Aedes* and *Anopheles* mosquitoes eggs, we used slightly higher concentrations of gRNA and Cas9 protein (200 ng/µl sgRNA and 200 ng/µl Cas9) compared to these other species in these experiments. Embryo survival post-microinjection ranged from 64-82% and somatic mutagenesis rates (*i*.*e*., mosaic eyes, with an intermediate wildtype black and white knockout phenotype, Fig. 1B) were 37 - 57% for single gRNA injections (Table 1). Notably, coinjection with two or more sgRNAs targeting different gene regions of *w* including: a) *w*sgRNA-1 and *w*sgRNA-2, b) *w*sgRNA-1 and *w*sgRNA-3, c) *w*sgRNA-2 and d) *w*sgRNA-3, *w*sgRNA-1, *-*2 and *w*sgRNA-3) increased G_0_ mutagenesis efficiencies to 74%, 73%, 78% and 86%, respectively. These results indicate that by using increased concentrations of CRISPR/Cas9 components we can achieve high single and multi-target somatic mutagenesis rates in *Cx. quinquefasciatus*, which is key to efficient CRISPR-mediated genome engineering in *Cx. quinquefasciatus.*

**Table 1.** Summary of the injection and mutagenesis mediated by independent sgRNAs in *Culex quinquefasciatus*

| sgRNA | #injected | Survival |  |  | Somatic Mosaicism (G<sub>0</sub>) |  |  | Germine Mosaicism (G<sub>1</sub>) |
| --- | --- | --- | --- | --- | --- | --- | --- | --- |
|  |  | ♀ | ♂ | Total (%) | ♀ (%) | ♂ (%) | Total (%) | G<sub>1</sub> mutants (%) |
| wsgRNA-1 | 50 | 15 | 20 | 35 (70) | 7 (47) | 13 (65) | 20 (57) | 128 (69) |
| wsgRNA-2 | 50 | 9 | 32 | 41 (82) | 3 (33) | 12 (38) | 15 (37) | 51 (61) |
| wsgRNA-3 | 50 | 17 | 15 | 32 (64) | 7 (41) | 8 (53) | 15 (46) | 157 (72) |
| wsgRNA-1/wsgRNA-2 | 50 | 7 | 16 | 23 (46) | 5 (71) | 12 (75) | 17 (74) | 123 (79) |
| wsgRNA-1/wsgRNA-3 | 50 | 13 | 9 | 21 (45) | 10 (77) | 6 (67) | 16 (73) | 72 (81) |
| wsgRNA-1/wsgRNA-3 | 50 | 17 | 10 | 27 (54) | 13 (76) | 8 (80) | 21 (78) | 101 (85) |
| wsgRNA-1/wsgRNA-2/wsgRNA-3 | 50 | 11 | 10 | 21 (42) | 9 (82) | 9 (90) | 18 (86) | 149 (86) |

**Figure 1.**
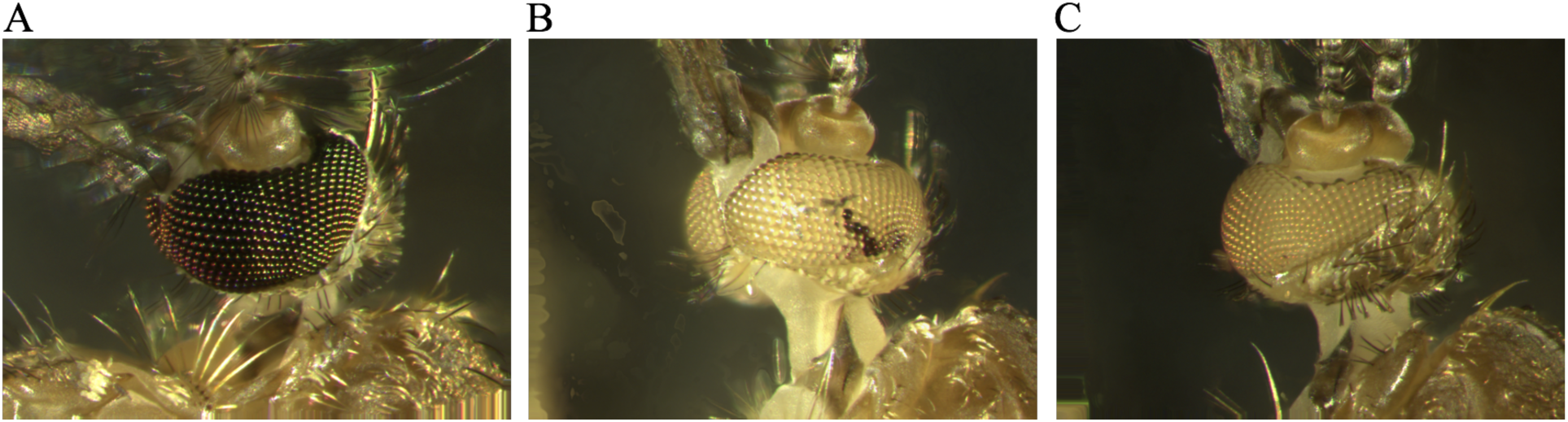
CRISPR/Cas9 efficiently generates heritable, site-specific mutations in *Culex quinquefasciatus*. (A) Representative image of wild-type *Cx. quinquefasciatus* adult eyes. (B) Representative G_0_ mosaic white-eyed mutant mosquito post-embryonic injection with a mixture of three unique sgRNAs targeting the *white* gene and the Cas9 endonuclease. (C) representative homozygous white-eyed mutant G_1_ mosquito generated by pairwise crossing mosaic G_0_ male and female mosquitoes. CRISPR, clustered regularly interspaced short palindromic repeats; sgRNA, small guide RNA.

**Figure 2.**
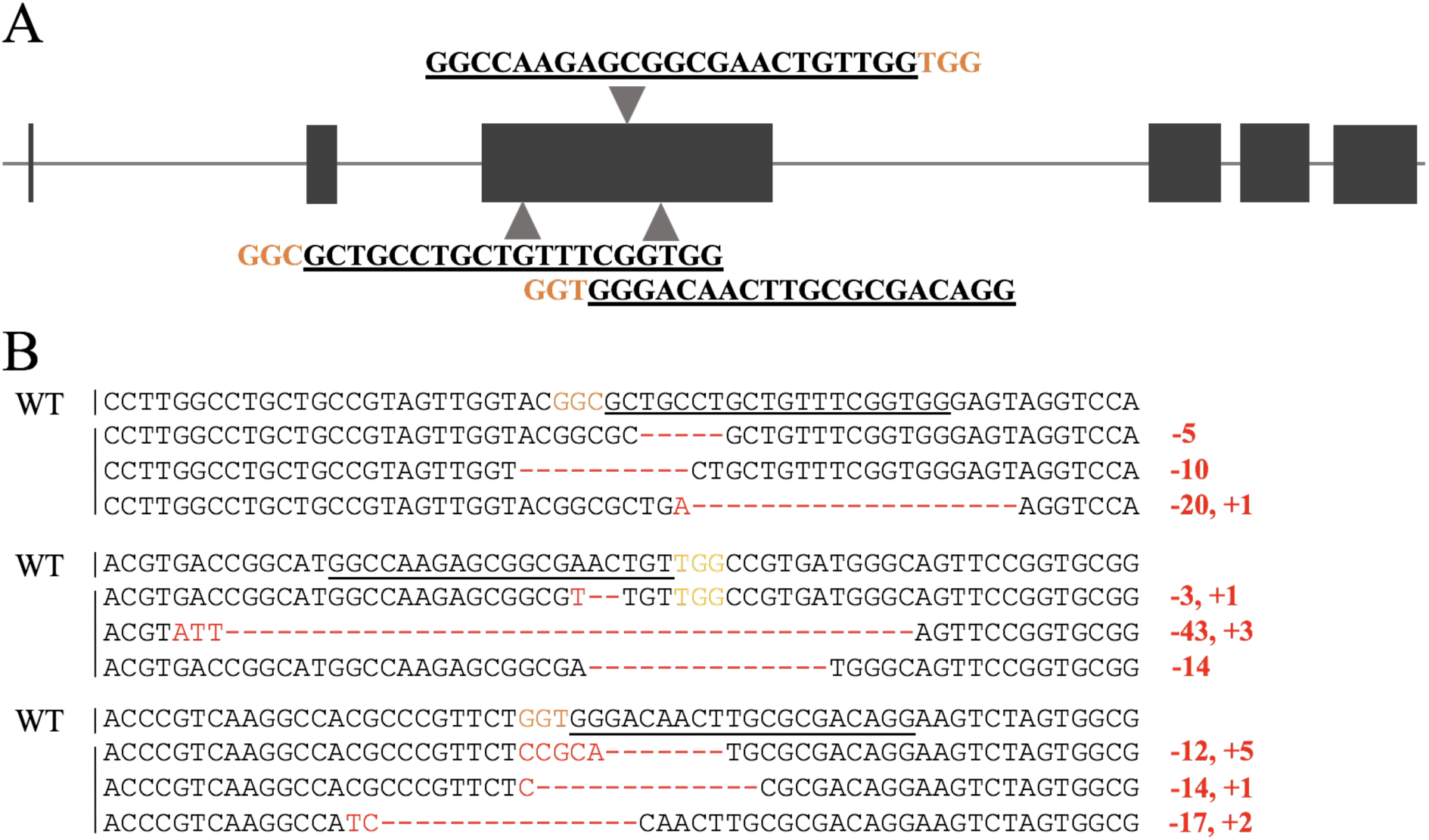
Mutagenesis of the *white* locus. (A) Schematic representation of the *white* locus with exons indicated as black boxes. Locations and sequences of the three sgRNA targets are indicated with the protospacer-adjacent motifs (PAM) highlighted in orange. (B) Genomic sequencing analysis of indels from individuals sequenced from the sgRNA injections. Top line represents wild-type (WT) sequence; PAM sequences (NGG) are indicated in yellow, and white gene disruptions resulting from insertions/deletions are indicated in red.

### Heritable mutations rates

Given the above promising results demonstrating efficient somatic editing, we next wanted to test for germline editing as this is required for heritable transmission of the engineered mutations. Germline mutation transmission efficiency from Cas9-directed genome engineering in *Cx. quinquefasciatus* was determined by intercrossing mosaic G_0_ males and females. The percentage of complete white eyed G_1_ progeny (Fig. 1C) in single target injections are shown in Table 1. G_1_ mutation rates increased to >79% when co-injected with multiple sgRNAs targeting different regions of the *w* gene. Deletion and insertion mutations in several independent mutant G2 lines were confirmed by sequencing the genomic DNA fragment containing the *w*sgRNA target sites (Fig. 2B). Some of the gene deletions were large, up to 43 bp (Fig. 2B). Homozygous viable healthy and fertile stocks were established for some of these *w* mutant lines indicating that this gene is not essential. Taken together, these results demonstrate that this method can generate germline mutations that can be inherited at a high rate and the efficiency of this inheritance is improved by synchronous injection of more than one sgRNA targeting different regions in the same gene.

## Discussion

The Itokawa et al. study is a great example of the important and high impact research that can be achieved with CRISPR/Cas9 mutagenesis in *Cx. quinquefasciatus*; however, this work did not provide the detail needed to effectively use this technology for future functional genomics studies in this vector, nor did it optimize the gRNA design, injection mixtures, egg handling and microinjection procedures (Itokawa et al. 2016). These details are particularly important as the unique biology of the *Culex* species eggs, which are laid in raft structures, require modifications from the standard microinjection procedures used for mosquitoes that lay single eggs, such as *Aedes* or *Anopheles* species. Proper separation, handling and injection of these egg rafts is not easy and took us many rounds of refinement to achieve the high survival and mutagenesis rates in this study. Earlier work that better describes *Cx. quinquefasciatus* microinjection procedures for *Hermes* transposable element-based transgenesis still had consistently lower survival rates (range 5 - 40% survival) (Allen et al. 2001) compared to our procedures (range 42-82% survival). Post injection survival rates were unreported in the 2016 study (Itokawa et al. 2016), so it is unclear whether they were more successful in their study, but as not enough details on methodology is provided, these results are unlikely to be recapitulated in another laboratory without significant troubleshooting.

Additionally, our work also outlines methods to optimize gRNA targeting and multiplexing to improve mutagenesis rates. Consequently, our design methods resulted in an increase in the mutagenesis rate of up to 26% for single gRNAs and up to a 40% for multiplexed gRNAs compared to the previous study (Itokawa et al. 2016) (Table 1). These improvements in survival and mutagenesis rates will make CRISPR-based technologies less laborious and more affordable to researchers and may even be generalizable enough to apply to other *Culex* species. Moreover, this is the first characterization of the *white* gene marker in *Cx. quinquefasciatus*, which is common phenotypic marker used in other mosquito species (Coates et al. 1997). This easily screenable phenotypic marker can be used to simplify the generation of new genetic tools and future functional genomic studies in this species. In the end, these procedures should provide the step by step instructions for the CRISPR/Cas9 directed mutagenesis of *Cx. quinquefasciatus* in any laboratory with the appropriate insectary facilities and therefore we think this work is an important step towards bridging the gaps in research for this important, but frequently overlooked, disease vector.

### Experimental Procedures

#### Mosquito strain and rearing

We used the *Cx. quinquefasciatus* wild-type S-strain (Li & Liu 2014) for these studies. Mosquitoes were maintained at the University of California, San Diego (UCSD). Mosquitoes were raised at 25.0 ± 1 °C with 30% humidity and a 12-hour light/dark cycle. A 20% sugar solution was provided daily. Females were offered a bovine blood meal using the Hemotek (model# PS5) blood feeding system.

#### sgRNA design and generation

gRNAs were designed to target both the sense and antisense strand of exon 3 of the *white* gene (CPIJ005542). Target and protospacer-adjacent motif (PAM) regions were selected using CHOPCHOPv2 (http://chopchop.cbu.uib.no/) and CRISPR Design (http://crispr.mit.edu/) (Xie et al. 2014). Linear, double-stranded DNA templates for sgRNAs were generated by performing template-free PCR with Q5 high-fidelity DNA polymerase (NEB) using the forward primer of each gRNA, and universal-sgRNAR. PCR conditions included an initial denaturation step of 98° for 30 seconds, followed by 35 cycles of 98° for 10 seconds, 58° for 10 sec, and 72° for 10 seconds, followed by a final extension at 72° for 2 min. PCR products were purified with magnetic beads using standard protocols. gRNAs were generated by *in vitro* transcription (AM1334; Life Technologies) using 300 ng purified DNA as template in an overnight reaction incubated at 37°C. MegaClear columns (AM1908; Life Technologies) were used to purify sgRNAs, which were then diluted to 1 μg/μl, aliquoted, and stored at 80°until use. Off-target sites of each sgRNA were identified based on the CHOPCHOPv2 software (Xie et al. 2014; Labun et al. 2016) and local sgRNACas9 package (Xie et al. 2014). All primer sequences are listed in Supplemental Material, Table S1 in <u>File S1</u>. Recombinant Cas9 protein from *Streptococcus pyogenes* was purchased from PNA Bio (CP01) and diluted to 1 µg/µl in nuclease-free water with 20% glycerol, and stored in aliquots at -80°.

#### Preparation of sgRNA/Cas9 mixtures for microinjection

The stock Cas9 protein solution was diluted with nuclease free water and mixed with the purified sgRNAs at various concentrations (20-320 ng/ul) in small 5-10ul aliquots to avoid excess freeze-thaw-cycles. These ready-to-inject mixtures were stored at -80C until needed. Mixtures were then thawed and maintained on ice while performing injections.

#### Preparation of needles for embryo microinjection

For effective penetration and microinjection into *Cx. quinquefasciatus* eggs, we experimented with several types of capillary glass needles with filament including quartz, aluminosilicate and borosilicate types. The quality of needles is critical for avoiding breakage/clogging during injection, embryo survival and transformation efficiency. For each of these glass types we developed effective protocols to pull these needles on different Sutter micropipette pullers (P1000, and P-2000) to enable the needles to have a desired hypodermic-like long tip that we found effective for *Cx. quinquefasciatus* embryo microinjection. The parameters (filament, velocity, delay, pull, pressure) for the different types of capillary glass needles are listed in the following table. While all three types of needles were effective for *Cx. quinquefasciatus* injections, we preferred the aluminosilicate capillary glass needles, because the quartz capillary glass needles were too expensive, and the borosilicate capillary glass needles were a bit too soft and clogged easily. All needles were beveled with a Sutter BV-10 beveler.

#### Embryo preparation and microinjection

Mixed sex pupae were allowed to eclose into a single (L24.5 × W24.5 × H24.5 cm) cage. Five days post emergence, mated females were offered a bovine blood meal using the Hemotek (model# PS5) blood feeding system. Then females were held for three to five days to allow oogenesis, after which oviposition cups with organically infused water were introduced into cages. Females were allowed to oviposit in the dark for 20-35 minutes. Fresh egg rafts (white color) were transferred from the cup to a wet filter paper with paintbrush. Single eggs were carefully separated for the egg raft under a dissecting scope with forceps. Single embryos were then aligned and a coverslip with double-sided sticky tape was used to secure the eggs. The eggs were then covered with halocarbon oil.

An Eppendorf Femtojet for the microinjections, which were performed under a compound microscope at 100× magnification. Needles were filled with 2 µls of injection mix and each egg was injected with the injection mixture at approximately 10% of the egg volume. We injected approximately 20 eggs per round and then carefully remove the halocarbon oil with a paintbrush. Eggs were then rinsed in ddH_2_0 and monitored for daily for hatching over the next week.

#### Mutation screens

The phenotype of G_0_ and G_1_ mosquitoes was assessed and photographed under a Leica M165 FC stereomicroscope. To molecularly characterize CRISPR/Cas9-induced mutations, genomic DNA was extracted from a single mosquito with a DNeasy blood & tissue kit (QIAGEN) and target loci were amplified by PCR. For T7EI assays, 1 μl of T7EI (NEB) was added to 19 μl of PCR product, digested for 15 min at 37°, and visualized on a 2% agarose electrophoresis gel stained with ethidium bromide. To characterize mutations introduced during NHEJ or MMEJ, PCR products containing the sgRNA target site were amplified, cloned into TOPO TA vectors (Life Technologies), purified, and Sanger sequenced at the source bioscience (https://www.sourcebioscience.com/).

## Data availability

Genomic DNA from mosquito strains produced here will be made available upon request.

## Acknowledgements

This work was supported in part by UCSD startup funds directed to O.S.A.

## Author Contributions

O.S.A, M.L and T.L. conceived and designed the experiments. M.L. and T.L. performed all molecular and genetic experiments. All authors contributed to the writing, analyzed the data, and approved the final manuscript.

---

**Table S1.**
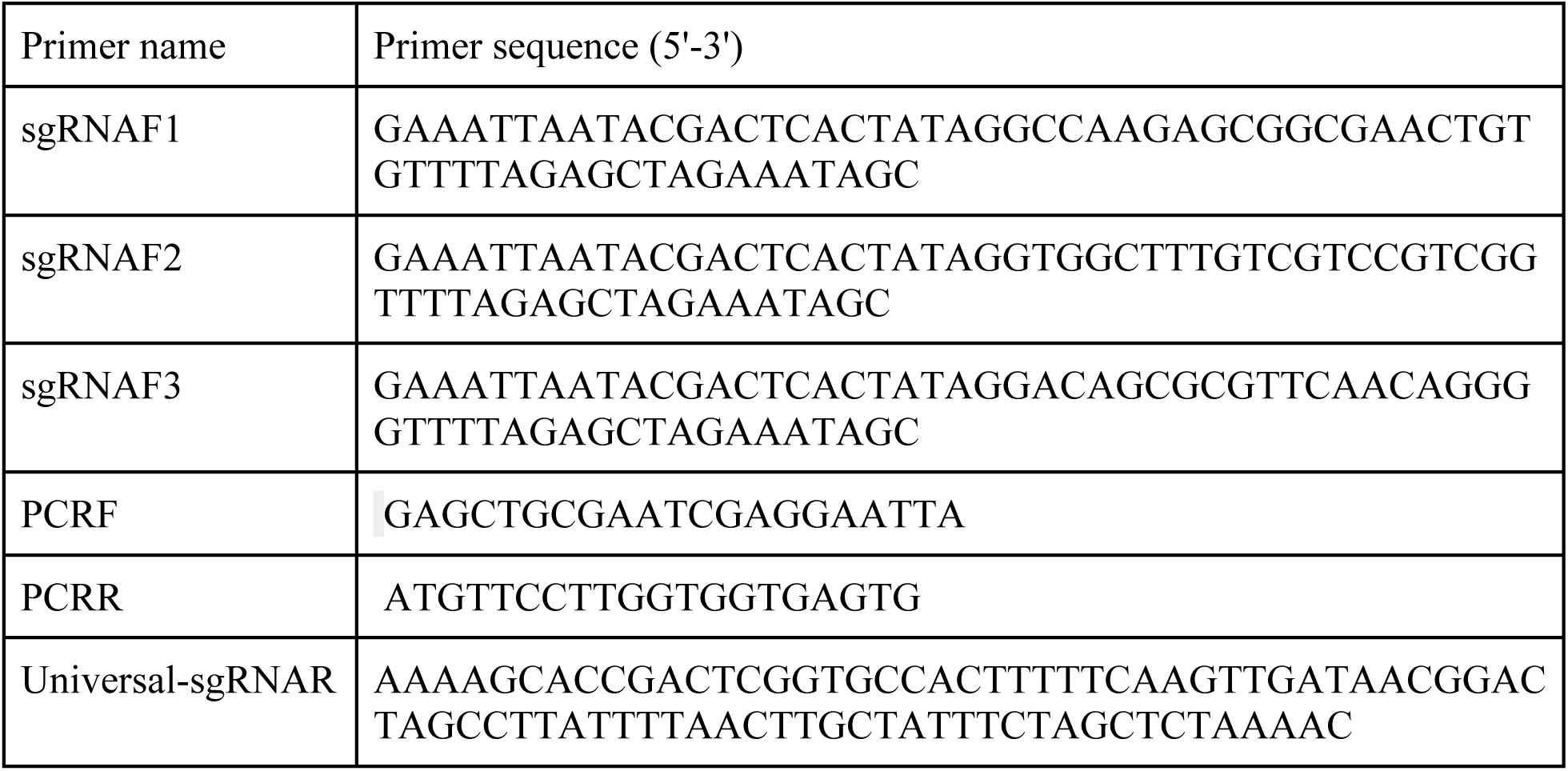
Primer sequences used in this study

**Table S2:** Optimized needle settings to limit needle breakage and clogging and to improve survival and transgenesis rates

| Capillary glass needle type | Sutter Needle Puller mode | Heat | Filament | Velocity | Delay | Pull | Pressure |
| --- | --- | --- | --- | --- | --- | --- | --- |
| Quartz | P-2000 | 750 | 4 | 40 | 180 | 135 | - |
| Aluminosilicate | P-1000 | 605 | - | 130 | 80 | 70 | 500 |
| Borosilicate | P-1000 | 450 | - | 90 | 80 | 70 | 500 |

